# Efficient indexing and querying of annotations in a pangenome graph

**DOI:** 10.1101/2024.10.12.618009

**Authors:** Adam M. Novak, Dickson Chung, Glenn Hickey, Sarah Djebali, Toshiyuki T. Yokoyama, Erik Garrison, Giuseppe Narzisi, Benedict Paten, Jean Monlong

## Abstract

The current reference genome is the backbone of diverse and rich annotations. Simple text formats, like VCF or BED, have been widely adopted and helped the critical exchange of genomic information. There is a dire need for tools and formats enabling pangenomic annotation to facilitate such enrichment of pangenomic references. The Graph Alignment Format (GAF) is a text format, tab-delimited like BED/VCF files, which was proposed to represent alignments. GAF could also be used to store paths representing annotations in a pangenome graph, but there are no tools to index and query them efficiently.

Here, we present extensions to vg and HTSlib that provide efficient sorting, indexing, and querying for GAF files. With this approach, annotations overlapping a subgraph can be extracted quickly. Paths are sorted based on the IDs of traversed nodes, compressed with BGZIP, and indexed with HTSlib/tabix via our extensions for the GAF format. Compared to the binary GAM format, GAF files are easier to edit or inspect because they are plain text, and we show that they are twice as fast to sort and half as large on disk. In addition, we updated `vg annotate`, which takes BED or GFF3 annotation files relative to linear sequences and projects them into the pangenome. It can now produce GAF files representing these annotations’ paths through the pangenome.

We showcase these new tools on several applications. We projected annotations for all Human Pangenome Reference Consortium Year 1 haplotypes, including genes, segmental duplications, tandem repeats and repeats annotations, into the Minigraph-Cactus pangenome (GRCh38-based v1.1). We also projected known variants from the GWAS Catalog and expression QTLs from the GTEx project into the pangenome. Finally, we reanalyzed ATAC-seq data from ENCODE to demonstrate what a coverage track could look like in a pangenome graph. These rich annotations can be quickly queried with vg and visualized using existing tools like the Sequence Tube Map or Bandage.

## Introduction

The current reference genome is the foundation enabling rich and diverse annotations. It is also used as a reference to map sequencing reads. Large annotation databases of functional elements, known variant information, and genomic elements use the coordinate system it provides. The organization and visualization of these annotations has been central to understanding and sharing results from genomic studies. In practice, annotations about a genome are typically saved separately from the genome itself, in formatted text files that can be compressed and indexed for fast query (e.g. VCF, BED, GFF). Under the hood, the HTSlib library supports the indexing of the most used file formats[1]. These annotation files are easy to write and to load in software like IGV[2] or websites like the UCSC Genome Browser[3].

With improved genome sequencing technologies, more high-quality genomes can be produced and combined into pangenomes. A pangenome represents multiple genomes, often as a graph describing adjacencies (edges) between the sequences (nodes) in the genomes. Tools to work with such pangenomes are still in their infancy, although there are now several applications where they show improved performance over traditional approaches. In particular, sequencing data analysis benefits from using pangenomes as a reference for read mapping[4,5], variant calling[6], peak calling[7,8], or transcript expression quantification[9]. Hence, we are now manipulating genomic objects, like sequencing reads, epigenetic regions, or genomic variants, in the pangenome graph space. Currently, those results are typically flattened or “surjected” onto the linear reference genome. As with the linear genome reference, organizing and visualizing genomic annotation in the pangenome will be essential. User-friendly querying and visualization options are required to adopt the pangenomic paradigm.

Several interactive visualization tools for pangenomes exist, but mainly focus on representing the graph topology, with tool-specific approaches to integrate additional information layers. We provide a short review of several of them here:

- **Bandage-NG**[10], a derivative of Bandage[11], can interactively visualize assembly graphs, and scales to pangenomes up to hundreds of thousands of nodes on modern computers.
- **GfaViz** is another interactive sequence graph visualization tool that supports version 2 of the Graphical Fragment Assembly (GFA) format and its new features[12]. In particular, it can represent the different paths annotated in the GFA file. GfaViz cannot load additional annotation files: the annotations, written for example as GFA *paths*, must be included in the main pangenome file to be accessible during exploration.
- **The Sequence Tube Map** visualizes the pangenome and sequencing reads aligned to it, allowing the user to query specific regions[13]. Prior to the present work, the application supported only the binary GAM format[4] for displaying additional information on the graph. It has some support for BED files, but only for inputting regions to display: BED-format annotations cannot be displayed.
- **MoMI-G**, for Modular Multi-scale Integrated Genome Graph Browser, focuses on visualizing structural variants[14]. Variants are interactively selected and represented with a pangenome view as a custom Sequence Tube Map representation. Supporting reads, and genome annotations from files in BED/GFF/WIG format, can also be included in that representation. Only the reference genome path can be annotated.
- **Panache** is another pangenomic tool specializing in gene-centric visualization of a linearized view of the pangenome, where blocks of homologous sequences are represented side by side[15]. The blocks are interactively explored in a web-based application to analyze their gene content and their absence/presence in the different genomes. Panache is notably used in the Banana Genome Hub[16].

In summary, while some visualization tools exist, there is no consensus on the best way to provide additional annotation layers to their graph representations.

Several other options exist to display static representations of a pangenome graph (or subgraph). The `vg` toolkit can output a pangenome and alignments in the DOT format. Paths can be displayed too, but the image can become hard to read when many paths are included or in large graphs. The `odgi` toolkit offers visualizations that scale better to larger graphs and more annotations[17]. The pangenome is linearized using a 1D layout[18], and annotations are displayed on top of the graph as an array/heatmap. Notably, `odgi` implements two options to add annotations from external BED files: one to convert annotations to a CSV file to help color nodes in Bandage, and another that injects BED records into the graph as new embedded paths that can appear in `odgi` ‘s visualizations. One limitation to this approach is that embedded paths are forced to cover entire nodes in the pangenome data model used by `odgi, vg`, and the GFA format. There is no way to provide an *offset* or *length* value to specify the base at which a path must start or end.

There is a dire need for a format supporting annotations in the pangenome that is easy to create, index, and query. Noting the success of the BED, GFF, or VCF formats, we see a critical need to exchange (pan)genomic information and allow tools to access additional information in the pangenomic space. The Graph Alignment Format (GAF) text format, proposed to represent alignments[19], could also describe annotations in a pangenome graph. However, the lack of techniques to compress, index, and query it limits its adoption for larger-scale annotation sets. Here, we present new features of the `vg` and HTSlib ecosystem that provide efficient indexing and querying for pangenomic annotations represented as paths in the GAF format. We illustrate its values in several applications: projecting gene and repeat annotations into the pangenome and visualizing them, summarizing open chromatin from epigenomic sequencing datasets, and positioning known variants in the pangenome.

## Methods

### Indexing paths in GAF files

GAF records are sorted by node ID intervals, so the sorting and indexing algorithm is most efficient. It makes sense when node IDs are integers, sorted based on the topology of the pangenome graph. This is the case for pangenomes constructed by minigraph-cactus[20], PGGB[21], or `vg construct` [4]. Otherwise, the pangenome graph can be *sorted*, by changing the node IDs, using `vg` or `odgi` [17]. If the node IDs are sorted integers, a short path through the graph should only traverse nodes with IDs in a small interval (see Figure 1A). The rationale behind our GAF sorting and indexing approach is to work with those intervals. Hence, to sort a GAF file, each path is first parsed to extract the minimum and maximum node IDs (see Figure 1B). The paths are sorted first by their minimum ID, and then by their maximum ID. Nodes are assumed to be short enough that within-node indexing is not required. This is similar to the approach used to sort BED or VCF files on linear genomes: they are sorted by sequence name, start position, and end position.

**Figure 1:**
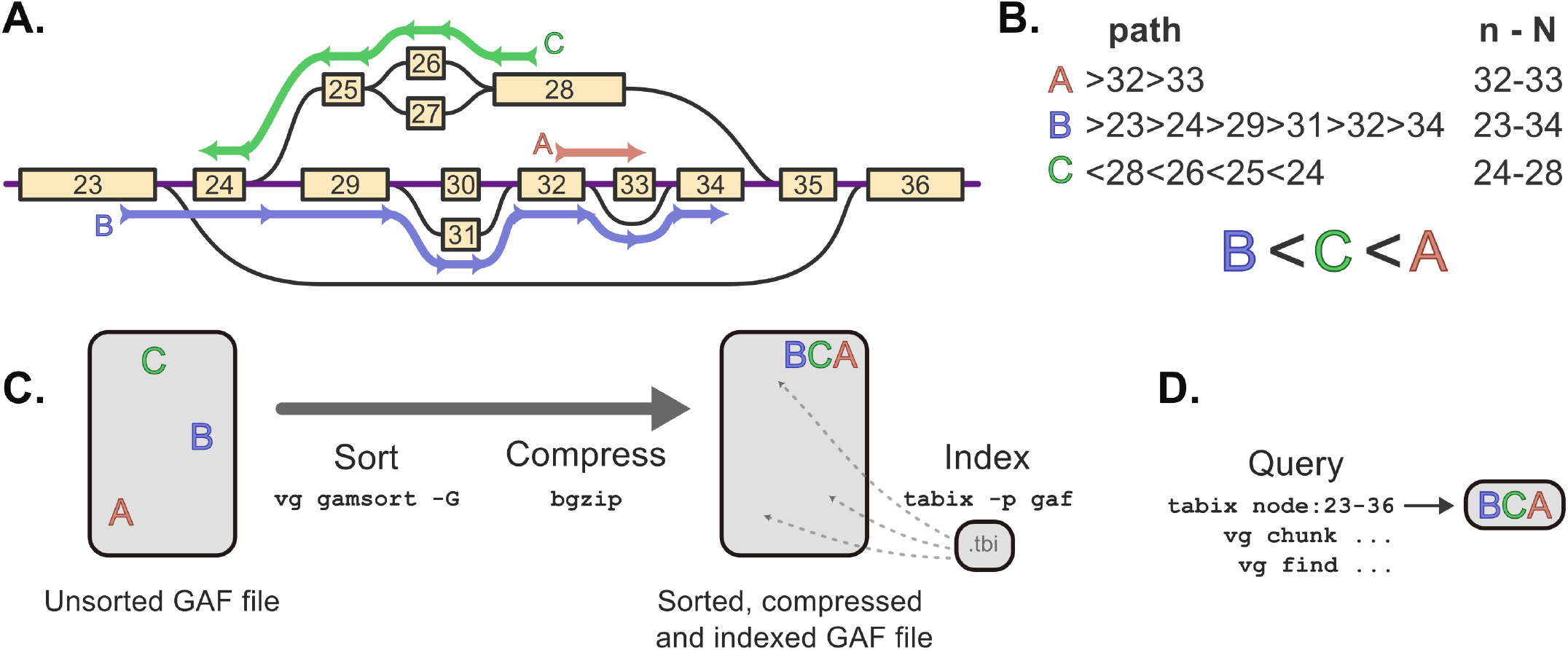
Path sorting and indexing using vg and HTSlib/tabix. **A**. A region of pangenome is represented with nodes (containing sequences) in yellow and edges in black. The node IDs are topologically sorted integers ranging from 23 to 36. Three paths are highlighted in red, blue and green. **B**. The three paths are written with the GAF syntax, specifying the traversed nodes’ orientations (< / >) and IDs. For each path, the node range *n-N*, between the minimum and maximum node IDs, is used for sorting the path. **C**. Overview of the workflow: Sort a GAF file using `vg gamsort`, compress it with `bgzip` and index using `tabix`. **D**. The small .*tbi* index file helps query slices of the GAF file quickly, and is usable with `tabix`, or `vg` subcommands like `find` or `chunk`.

We added a GAF sorting feature to the `vg` toolkit’s `gamsort` subcommand, available in vg version 1.56 and later (see Figure 1C). It first sorts chunks of GAF records (e.g. reads), to avoid holding the entire set in memory. The sorted chunks are then merged into the final sorted GAF file. A sorted GAF file can be compressed in the BGZF format (or *bgzipped*) using `bgzip`. The BGZF format is a backwards-compatible extension of the standard gzip format that compresses a file by block to facilitate random access. It is widely used to compress several bioinformatics file types, including VCF, BED, and BAM files, and is implemented in HTSlib[1].

We extended HTSlib[1] to allow indexing bgzipped GAF files. Similar to other tab-separated file like VCF or BED, a `gaf` preset was added to `tabix`. For BED or VCF, `tabix` extracts the interval information from the columns with the sequence name and genomic position[22]. In the new `gaf` preset, it instead parses the path information in the GAF file to extract the minimum and maximum node IDs. The indexing is then based on this interval, as for the sorting described above.

We tested the GAF sorting and indexing performance on 30X coverage Illumina short reads from HG002. The reads were downloaded from https://s3-us-west-2.amazonaws.com/human-pangenomics/NHGRI_UCSC_panel/HG002/hpp_HG002_NA24385_son_v1/ILMN/downsampled/. We mapped them using `vg giraffe` [5] on a personalized pangenome[23] derived from the HPRC v1.1 Minigraph-Cactus pangenome. The aligned reads were first stored in GAM, to allow comparing file size and sorting runtimes against GAF. This unsorted GAM file was sorted using `vg gamsort`. In parallel, the unsorted GAM was also converted to a GAF file using `vg convert`, compressed with plain gzip, and sorted using `vg gamsort` with the new GAF sorting mode described above, with BGZF compression from `bgzip`. We compared the file sizes and sorting runtimes between both approaches. The query time was also measured on ten thousand regions of the pangenome, specified as node ID ranges. These ranges were created by picking a random starting node position on the reference path and walking 50 steps along that path. The commands and details for this benchmarking are available in the `analysis/readsorting` folder of this paper’s repository (see Code and data availability).

### Querying GAF files

Instead of querying a sequence name and genomic position, we can query a node interval (see Figure 1D). In HTSlib, `tabix` was modified to disregard any provided sequence name when querying intervals for a GAF file, and to interpret the values typically used for the *start* and *end* position of genomic coordinates as a node ID interval.

Commands to query slices of the pangenome in `vg` were also updated. The `find` and `chunk` sub-commands now use the updated HTSlib library to extract the appropriate GAF records. Internally, these commands identify an initial subgraph, which might correspond to, for example, a genomic interval on the reference path provided by the user. This subgraph is then extended with additional *context*, allowing a linear-reference-based query to include non-reference regions of the graph. The user controls the amount of context added. Finally, the paths (historically reads) overlapping the subgraph nodes are extracted. This last step was updated to extract paths from an indexed GAF file using HTSlib. It is now possible to extract a slice of an indexed GAF file based on node intervals, coordinates on a reference path, a list of coordinates from a BED file, or a user-provided subgraph, as described in the User Guide in the `analysis` folder of this paper’s repository (see Code and data availability).

### Projecting annotations into a pangenome

A pangenome represents multiple genomes for which annotations might be available. We describe an approach to project annotations relative to a genome into a pangenome. We updated the `annotate` subcommand from the `vg` toolkit to project regions represented in BED or GFF files into pangenomes and tested it on GBZ-format pangenomes from the HPRC. The annotation source genome must be marked as a *reference* in the pangenome. However, any genome can be quickly (about a minute in our environment) set to be a reference using the `vg gbwt` subcommand. Then, `vg annotate` traces out the path in the pangenome graph taken by each input annotation. The path, represented as an *alignment* record, is then written in GAM or GAF format. Sometimes, an annotation’s source path might be broken into multiple disjoint parts, because some regions of the individual assemblies are clipped out of the pangenome. Newly assembled centromeres that challenge current alignment tools, and regions whose alignments would introduce suspiciously large structural variants, are often excluded when pangenomes are constructed from assembled sequences. Projected annotations are thus also broken up by `vg annotate` as needed when they overlap with breakpoints in the pangenome’s representation of their source path. The name of the annotation record in the output file is taken from the BED file’s *name* column, or the *Name* and *ID* GFF fields.

We tested this approach by projecting the gene annotations, repeat annotations, and segmental duplication annotations from each of the 88 assembled haplotypes from the HPRC Year 1 data release into version 1.1 of the GRCh38-based HPRC Minigraph-Cactus pangenome. The annotations were downloaded from the HPRC public repository at https://github.com/human-pangenomics/HPP_Year1_Assemblies/tree/main/annotation_index. The gene annotations from CAT[24] (GFF files) were downloaded using the URLs in the Year1_assemblies_v2_genbank_CAT_genes.index file, the repeat annotations from RepeatMasker (BED files) from Year1_assemblies_v2_genbank_Repeat_Masker.index, segmental duplications (BED files) from Year1_assemblies_v2_genbank_Seg_Dups.index, and simple repeats from Year1_assemblies_v2_genbank_TRF.index. A helper script was implemented to prepare the BED files with informative record names. For example, for repeats from the RepeatMasker annotation, we name each element by its repeat class and repeat family. The projection of those annotations for each haplotype into the pangenome was automated with a Snakemake workflow available in the `analysis/annotate` folder of this paper’s repository (see Code and data availability).

### Coverage track from mapped reads

Functional genomics datasets are often visualized as a coverage track. High coverage in a region might suggest a strong transcription factor binding site or regulatory region.

We implemented an approach to summarize the coverage of reads across the pangenome into paths along which coverage is consistent (see Figure 2). The coverage in every node is first binned into a few coverage classes, for example representing low, medium and high coverage. By default, we use 1, 5, and 30 reads as coverage breakpoints to save three bins: 1-5, 5-30, 30+. Regions with no coverage are not saved. Once the coverage is binned, we extend greedily to connected nodes and bins if in the same coverage class. This extension step produces a path through the pangenome with consistent coverage. The paths are written in the GAF format and the coverage class and the average coverage across the path are recorded.

**Figure 2:**
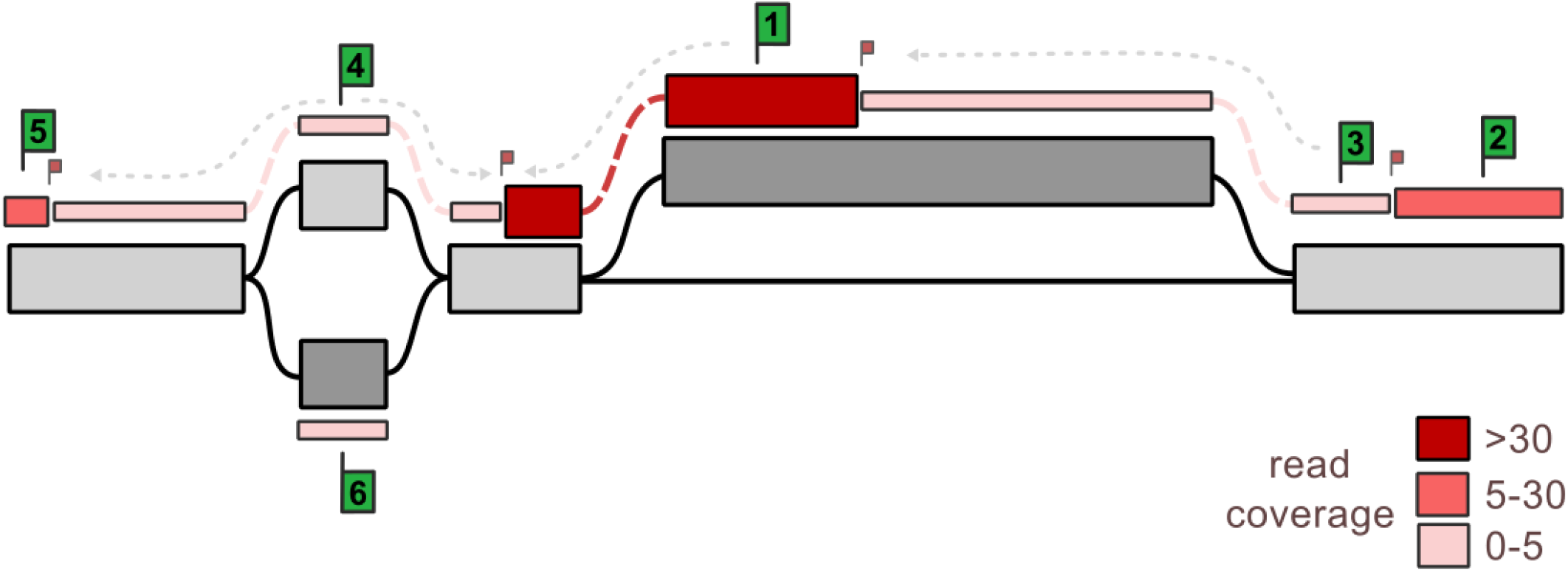
Read coverage bin-and-extend approach to produce coarse-grained coverage tracks. In each node (*grey rectangles*), the read coverage at each base is binned using user-defined coverage bins, creating node regions of similar coverage, also called bins (*colored rectangles*). Starting bins (*large green flags*) are selected one at a time, and those bins are extended in both directions until reaching a different-coverage bin (*small red flag*).

This algorithm was implemented in Python and uses the *libbdsg* module[25] to query the pangenome. It is made available in the public repository of this study in the `analysis/encode` folder of this paper’s repository (see Code and data availability).

We tested this approach on ATAC-seq datasets from ENCODE[26] on seven tissues: breast epithelium (*ENCFF210QPG*), gastrocnemius medialis (*ENCFF968KAQ*), gastroesophageal sphincter (*ENCFF553SEZ*), Peyer’s patch (*ENCFF272PIN*), sigmoid colon (*ENCFF775DFR*), spleen (*ENCFF391IHY*), thyroid gland (*ENCFF579WLS*). Paired-end short Illumina HiSeq 4000 reads were downloaded from ENCODE and mapped using giraffe[5] on the HPRC v1.1 Minigraph-Cactus pangenome. The output to `vg pack` (with -`Q 1` to keep reads with mapping quality of at least 1) was piped to the script implementing the coverage track computation described above.

### Annotating known variants

We implemented an approach to find variants from public databases in the pangenome, producing a GAF file with a pangenome path for each variant. To accomplish this, we match input variants against the VCF representation of the pangenome, produced by running `vg deconstruct` on the pangenome file. This VCF file contains all variants and their paths through the pangenome in the *AT INFO* field. We look for variants in the pangenome VCF that overlap each input variant. When an overlapping variant is found, we extract one of two paths from its *AT* field. If the input variant is found to be present in the pangenome VCF, we extract the path for the alternate allele. If the alternate allele of the input variant is not present, but there is still a variant at this position in the pangenome, we extract the path for its reference allele instead. Variants that are not matched in the pangenome are skipped. This algorithm was implemented in Python and uses the *libbdsg* module[25] internally to get the node size information necessary to write GAF records.

We tested this approach on the GWAS Catalog[27] and GTEX expression QTLs[28], and matched them against the HPRC Minigraph-Cactus v1.1 pangenome. The GWAS Catalog was downloaded from the UCSC Genome Browser’s `gwasCatalog` table[3]. The eQTLs from GTEX v8 were downloaded from https://storage.googleapis.com/adult-gtex/bulk-qtl/v8/single-tissue-cis-qtl/GTEx_Analysis_v8_eQTL.tar. The files defining associations between variant/gene pairs were parsed for each tissue. As before, the output annotations in GAF were bgzipped, sorted, and indexed. We collected statistics about the resulting annotations, and visualized database representation of variants using the Sequence Tube Map.

We implemented another, more straightforward approach, to annotate variants that were genotyped using the pangenome. Here, we simply convert a VCF that contains allele traversal information (*AT* field) to a GAF file representing the alternate allele. We test this approach by genotyping HG002 from the short-reads Illumina Novaseq reads aligned to the pangenome for the GAF sorting experiment (see Indexing paths in GAF files). Variants were genotyped from the aligned reads using `vg call`. Notably, `vg call` can also output genotypes directly in GAF format with the `-G` parameter.

The scripts and pipeline to annotate variants is available in the public repository of this study in the `analysis/variants` folder of this paper’s repository (see Code and data availability).

### Visualization in the Sequence Tube Map

The Sequence Tube Map was developed to explore a pangenome graph interactively, with haplotypes traversing it, and reads mapping to it[13]. It internally calls *vg* to extract the relevant slice of the pangenome graph and reads, which previously could only be in the GAM format. We updated the Sequence Tube Map to accept GAF files sorted, compressed, and indexed as in Indexing paths in GAF files. We have also extended it to use an element’s opacity to represent an integer score, like a mapping quality or read coverage. The variable transparency helps highlight regions of high coverage when visualizing coverage tracks from the ENCODE project (see Coverage of seven functional datasets from ENCODE and Figure 4D).

### Visualization in Bandage-NG

Bandage-NG[10], a tool derived from Bandage[11], can read GAF files and visualize its paths through the input graph by coloring the nodes. Of note is that the offset positions are not considered, which means that nodes are colored entirely, even if the GAF record only overlaps part of it. We implemented a wrapper script to facilitate the preprocessing of a subgraph to visualize with Bandage-NG. It starts by extracting the subgraph for a region of interest from a full pangenome using `vg chunk`. This graph is converted to GFA, ensuring the paths are written last for compatibility with Bandage-NG. The script also extracts annotations relevant to the region of interest from indexed GAF files. The extracted GAF files are merged and modified if necessary: path names are de-duplicated, and paths are trimmed when traversing nodes absent from the extracted subgraph. These modifications ensure that the final GAF file can be loaded in Bandage-NG. The outputs of this script can be opened with Bandage-NG for interactive exploration. In particular, the *Find path* feature can find nodes corresponding to specific paths, either haplotypes or annotations in this pre-processed GFA file. The user can select a *path* and color its nodes. The helper script and a tutorial are available at the `analysis/visualization` folder of this paper’s repository (see Code and data availability).

## Results

### Sorting and indexing short sequencing reads

Read sorting (see Indexing paths in GAF files) was tested on about 30X coverage of Illumina HiSeq paired-end 150bp reads for the HG002 sample. The plain gzipped GAF file with about 682 million reads was sorted with `vg gamsort` and compressed with `bgzip` in 6h32 using 6.47 hours of CPU time and about 1.9 GiB of memory (Table 1). Indexing the sorted GAF with `tabix` took 18.6 minutes using 17.4 minutes of CPU time. We compared that approach with the existing read sorting implementation in `vg`, which operates on files in the Protobuf-based GAM format[4]. Sorting a GAM with the same reads took 11h47 using 11h38 of CPU time and about 6.1 GiB of memory.

**Table 1.** Resources used to sort short sequencing reads for a 30x coverage Illumina human dataset.

| Format | Time (H:M:S) | Max. memory used (MiB) | File size (GiB) |
| --- | --- | --- | --- |
| GAM | 11:46:58 | 6,236.60 | 108 |
| GAF | 6:50:28 | 1,904.83 | 52 |

In addition to being about twice as fast to sort, reads written in the GAF format (and bgzipped) also take about half the disk space (52 GiB vs 108 GiB). The main reason for this reduced space is that GAF does not save the complete read sequence, only the path through the pangenome and edits to reconstruct it. GAM files produced by `vg giraffe` can also save additional information as annotations, like the mapping time of each read, which are currently not kept when converting to the GAF format. Overall, the bgzipped GAF files are half as small and twice as fast to sort for short sequencing reads.

Once a GAF is indexed, extracting a slice of reads works like extracting a slice of an indexed BAM/VCF/BED file for a genomic region, and is more efficient than extracting a slice from an indexed GAM file. For example, extracting reads for ten thousand random regions in the pangenome took about 0.066 seconds per region to retrieve an average of 1707 reads. For comparison, the same extraction took an average of 0.816 seconds per region using the GAM format.

### Annotation of a human pangenome

To showcase our annotation projection implementation, we projected annotations for all HPRC haplotypes into the HPRC pangenome (see Projecting annotations into a pangenome). This included genes, segmental duplications, tandem repeats, and repeat annotations. `vg annotate` was able to project ∼4M gene annotations into the pangenome in ∼11 minutes on two cores using 19.9 minutes of CPU time and 21 GiB of RAM. It projected ∼5.5M repeats from RepeatMasker in ∼2.9 minutes on two cores using 4.9 minutes of CPU time. These measures include projection with `vg annotate`, decompression of the input gzipped GAF, and compression of the output GAF with `bgzip`. We could quickly query these rich annotations with `vg`, and visualize them using tools like the Sequence Tube Map or Bandage-NG. Using Bandage-NG, we prepared a visualization illustrating a mobile element insertion (Figure 3). We also examined a gene annotation using the Sequence Tube Map (Figure 4A).

**Figure 3:**
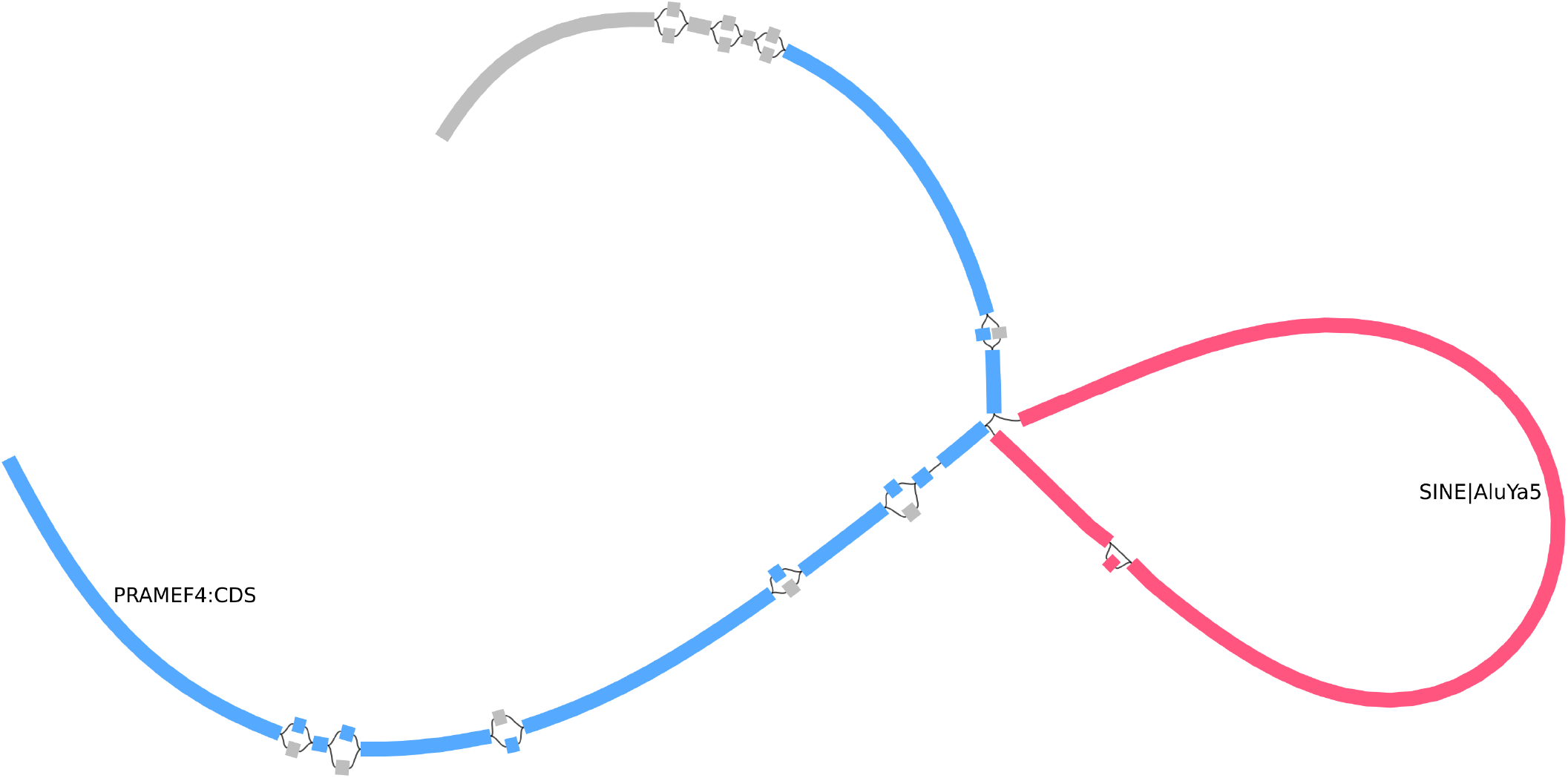
Visualization examples with Bandage-NG. Example of an AluYa5 transposon insertion (*red*) within the coding sequence of the *PRAMEF4* gene (*blue*). Both annotations were initially produced at the haplotype level by the Human Pangenome Reference Consortium. We projected them into the pangenome, indexed them, and queried a small region to visualize with Bandage-NG. The nodes were colored based on those annotations and loaded as paths by Bandage-NG.

**Figure 4:**
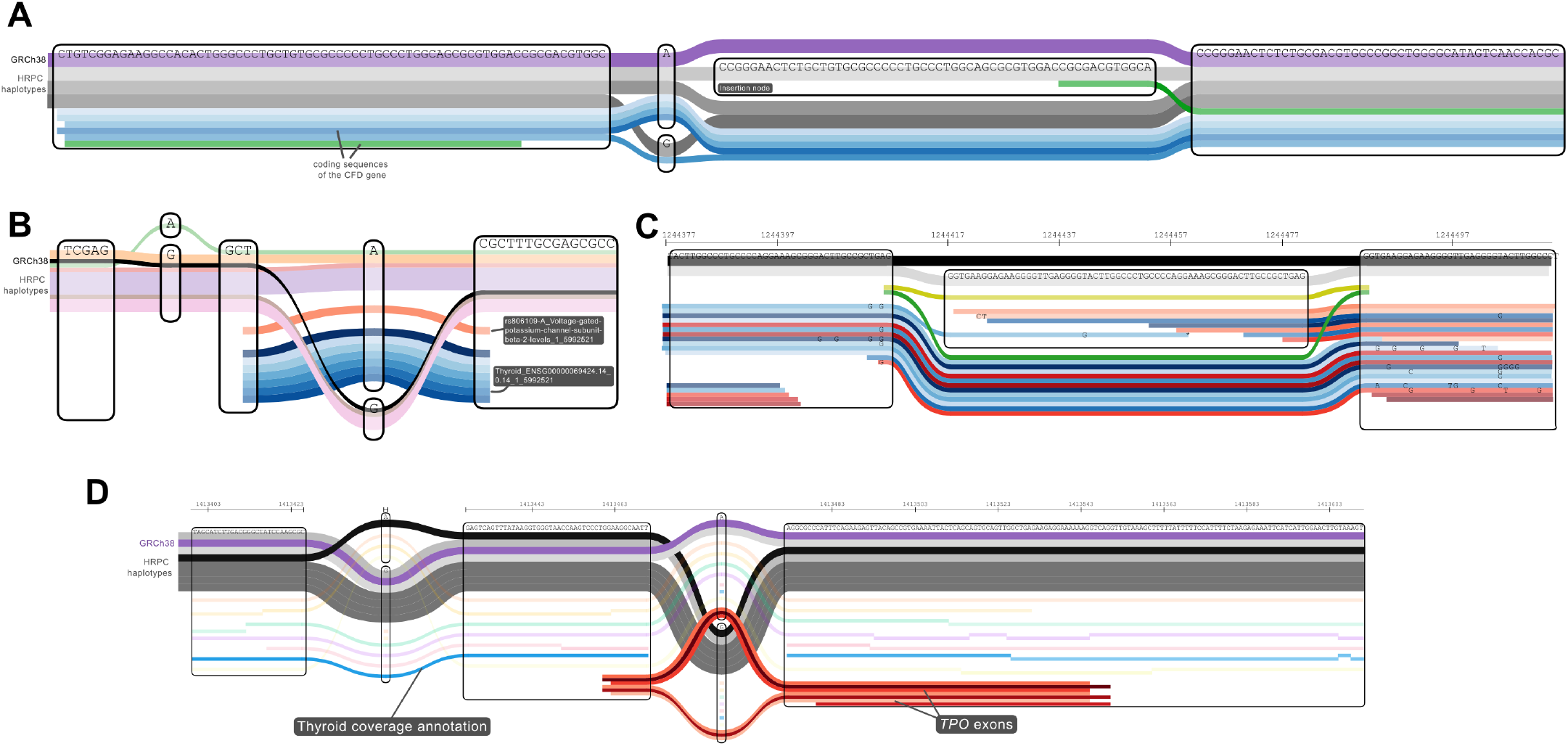
Visualization examples with the Sequence Tube Map. **A** Projection of the HPRC gene annotation into the pangenome, here visualizing a coding sequence (CDS) of the *CFD* gene. There is an insertion (see *insertion node* in the middle), that the reference path (*violet* on top) skips. Most CDS annotations are similar (*blue* paths), but the CSD from the haplotype with the insertion is split into two parts, highlighted in *green*. **B**. Projecting known variants into the pangenome. *black*: Reference path (GRCh38), *pale colors*: other human haplotypes, *reds*: GWAS Catalog, *blues*: eQTLs across tissues (GTEx). **C**. Aligned reads and genotype calls. *yellow*/*green*: annotation paths from `vg call` genotypes. *reds*/*blues*: short sequencing reads. **D**. ATAC-seq coverage track for seven tissues from the ENCODE project. The reference path (*violet*) and other HPRC haplotypes (*greys*) are shown on top and traverse two variation sites. ATAC-seq coverage annotations for the different tissues are shown in different *colors* with different opacities representing coverage level. The thyroid annotation (*blue*) is more opaque than others, suggesting that this region is more open in that tissue. The *red* annotations at the bottom show the position of exons from the *TPO* gene, a thyroid-specific gene.

We also matched and annotated variants from the GWAS Catalog[27] and expression QTLs from the GTEx catalog[28] across 49 tissues (on average 1.45 million variants per tissue), using the methods described in Annotating known variants. On average, 94% of variants were found in the HPRC pangenome. The variants files in GAF take only 907 MiB of space and can be queried rapidly for visualization in the Sequence Tube Map or Bandage-NG. This annotation is showcased in Figure 4B.

It is also straightforward to convert genotypes called on variants from the pangenome back to annotation paths. This is applicable when the pangenome is used as a source of variants for genotyping in new samples, as when using Pangenie or `vg call`. To illustrate this use case, we genotyped HG002 using short Illumina reads aligned to the HPRC pangenome with `vg giraffe` and `vg call` (see Annotating known variants). The predicted genotypes were converted to GAF and indexed. They could be visualized using the Sequence Tube Map, along with the aligned reads. This type of annotation can help a developer explore variant calls and aligned reads. An example is shown in Figure 4C. On the traditional reference genome, the equivalent would be to load both reads and variants in IGV[2].

### Coverage of seven functional datasets from ENCODE

We aligned ENCODE ATAC-seq datasets from 7 cell types[26] to the draft human pangenome to produce coverage tracks as indexed GAF files (see Coverage track from mapped reads). On average across cell types, there were about 475 thousand paths representing high read coverage, which were, on average, 2.6 nodes and 104.8 bases long. An average of 63 thousand paths with high ATAC-seq read coverage traversed three or more nodes (see Table 2). Because the graph node length used in non-variable regions is longer than the path length, these paths visiting three or more nodes are likely in regions of the pangenome with variation.

**Table 2.** High coverage tracks from seven functional datasets on the HPRC pangenome. For each sample, the table shows how many paths had a mean coverage of at least ten reads and how long they were.

| Dataset | Paths | Average bases | Average nodes | Traversing >2 nodes |
| --- | --- | --- | --- | --- |
| Breast epithelium | 570,155 | 109.80 | 2.68 | 65,095 (11.4%) |
| Gastrocnemius medialis | 342,830 | 94.47 | 2.43 | 53,595 (15.6%) |
| Gastroesophageal sphincter | 555,618 | 115.69 | 2.73 | 72,407 (13%) |
| Peyer's patch | 270,094 | 97.88 | 2.62 | 43,591 (16.1%) |
| Sigmoid colon | 678,380 | 115.10 | 2.69 | 82,906 (12.2%) |
| Spleen | 531,910 | 104.97 | 2.66 | 67,153 (12.6%) |
| Thyroid gland | 377,157 | 95.63 | 2.41 | 56,441 (15%) |

It took on average 7 CPU-hours to map the reads to the pangenome using VG Giraffe, and 2.8 CPU-hours to produce the coverage tracks. Sorting, compressing and indexing them took only 0.23 CPU-hours, on average. Table 3 compiles the runtime and memory usage for each step across all samples.

**Table 3.** Compute resources used for the analysis of the functional datasets and production of the indexed coverage tracks. The *coverage track* step includes computing coverage profile on the pangenome using `vg pack`, making coverage tracks using a python script, and compressing the output GAF with `gzip`.

| Dataset | reads (M) | read mapping | coverage track | sorting + compressing + indexing |
| --- | --- | --- | --- | --- |
| Breast epithelium | 193.6 | 8.9 CPU-H (54 GiB) | 3 CPU-H (109 GiB) | 0.3 CPU-H (1 GiB) |
| Gastrocnemius medialis | 98.8 | 4.8 CPU-H (54 GiB) | 2.6 CPU-H (99 GiB) | 0.3 CPU-H (1 GiB) |
| Gastroesophageal sphincter | 168.5 | 7.3 CPU-H (54 GiB) | 3 CPU-H (108 GiB) | 0.2 CPU-H (1 GiB) |
| Peyer's patch | 145.3 | 8 CPU-H (54 GiB) | 3 CPU-H (104 GiB) | 0.2 CPU-H (1 GiB) |
| Sigmoid colon | 173.5 | 8.2 CPU-H (54 GiB) | 3 CPU-H (106 GiB) | 0.3 CPU-H (1 GiB) |
| Spleen | 157.2 | 7.6 CPU-H (54 GiB) | 3 CPU-H (104 GiB) | 0.2 CPU-H (1 GiB) |
| Thyroid gland | 91.4 | 4.6 CPU-H (54 GiB) | 2.5 CPU-H (94 GiB) | 0.1 CPU-H (1 GiB) |

Figure 4D shows an example of a region, visualized with our extended version of the Sequence Tube Map (see Visualization in the Sequence Tube Map), near an exon of the *TPO* gene. The exon annotations came from projecting the HPRC gene annotations described above into the pangenome. The coverage track for the ATAC-seq dataset shows that this region is only opened in the thyroid, which is consistent with the thyroid-specific role of *TPO*. By integrating these two external sources of annotations (gene annotation and ATAC-seq coverage), we can visualize them in the context of the genomic variation in the pangenome.

## Discussion

The tools and applications described here streamline the production of annotations in the GAF format, their indexing, and efficient querying.

First, these techniques will help integrate additional information into existing or future pangenomic tools. They will simplify integrating information like gene or repeat annotations, known variants, or, more generally, other functional annotations in pangenomic analyses. These analyses will be able to work with those annotations in light of underlying genomic variation described in the pangenome. Visualization is a specific example where the users typically want to include several layers of annotations, including some of their own design. Tools to visualize pangenomes will greatly benefit from a simple pipeline to create or load pangenomic annotations.

Second, this work is critical to allow the community to make, share, and reuse annotations in the pangenome space. The GAF format is already used by multiple independent tools, although mostly for read mapping. Thanks to our work, GAF files can now be indexed and queried efficiently, like BED, VCF, or GFF on a linear reference genome. This will be helpful for those working with GAF-format read mappings and enable the format to be used for large annotation sets. We showcase how annotations on a linear genome can be projected into the pangenome and stored in GAF format. We believe it could become the *de facto* format to represent annotations on the pangenome, thus accelerating the broader genomics field’s adoption of the new pangenomic paradigm.

The indexing scheme relies on integer node IDs, compacted relative to their position in the pangenome. If node IDs are not integers, it adds a layer of complexity for the user as they will need to convert them to integers and keep track of the translation between original and new pangenome.

Both `vg` and `odgi` offer ways to support this, but tool support for the resulting translation information is uneven and no standard format exists to store it.

Our implementation of GAF-based annotation is designed with short paths in mind, where the user wants to extract the full annotation paths for all annotations in a region. This is convenient when working with short reads, gene annotations, and most genomic repeats, for example. Larger genomic regions, like chromosome bands, large assembled contigs, or large segmental duplications, can still be represented and manipulated, but might be less practical to work with. The current implementation extracts all annotations overlapping a queried region. The extracted annotations are not clipped to the specified range, and no tooling support is provided for finding the part or parts of each annotation record that are relevant to the query. To address this use case, we plan to implement an extraction mode where output annotations are trimmed to keep only parts overlapping the queried subgraph.

The naïve metadata integration is another limitation. Indeed, the input annotation information from BED and GFF3 files is currently reduced to a single string label. In our applications, those labels contain, for example, a gene name and haplotype names, or a read coverage bin and average coverage value, all in one label. This was easy to implement and sufficient for now because no tools handle more advanced metadata organization. For many annotations, keeping the metadata better organized would be useful so that the user can access/use it within visualization tools. The different metadata fields could be saved using GAF tags that can be added at the end of each GAF record.

Our approach to converting annotations from a linear genome to the pangenome assumes we have annotations of the different haplotypes in the pangenome. There is still no clear solution to lift annotations from one reference/haplotype to other haplotypes in the pangenome; reanalysis/reannotation of each haplotype is generally required. The homology information embedded in the pangenome can propagate annotations from one haplotype to others more easily. This strategy is used by annotation tools like CAT[24], and improves the reanalysis of each haplotype from scratch. In the future, these techniques might help propagate other annotations across the pangenome more efficiently than by reanalyzing the raw data from scratch on each haplotype.

Our GAF-based approach also highlights the limitations of the existing tool support for GAF files. Some tools, like Bandage-NG or GfaViz, require manual pre-processing, such as extracting and post-processing a subgraph and annotations. The Sequence Tube Map can now handle indexed bgzipped GAF files, but the query time for large pangenomes remains long in practice. Integrating annotation metadata into its interface as a first-class data type will also require a significant amount of development. Overall, we stress the need for visualization tools to lay out and organize many paths through a pangenome efficiently.

We have shown that the GAF format, thanks to new tools for its efficient manipulation, could offer a path for the future of annotations in pangenome graphs. While it provides an important building block, it is clear that more needs to be done to make it a useful solution for the community.

## Code and data availability

`vg` is available at https://github.com/vgteam/vg. Bandage-NG is available at https://github.com/asl/BandageNG. The Sequence Tube Map is hosted at https://github.com/vgteam/sequenceTubemap. The modified version of the Sequence Tube Map used to make Figure 4 is available in the quay.io/jmonlong/sequencetubemap:vggafannot Docker container.

The analysis presented in this manuscript is documented in the repository at https://github.com/jmonlong/manu-vggafannot[29]. It contains scripts to prepare the annotation files, commands and automated pipelines to annotate them in the pangenome, and helper scripts to summarize the output files.

Annotation files produced for this work were deposited on Zenodo at https://doi.org/10.5281/zenodo.13904205 [30]. This includes haplotype annotations of the HPRC v1.1 pangenome (gene annotations, repeats from RepeatMasker, simple repeats), coverage tracks for the seven ENCODE ATAC-seq samples, eQTLs from GTEx and the GWAS Catalog.

## Acknowledgments

Integration of GAF in SequenceTubeMap was supported by the National Cancer Institute of the National Institutes of Health under Award Number U01CA253405. We would like to thank the ENCODE consortium and the laboratory of Michael Snyder for making the ATAC-seq datasets available.

## Author contribution statement

AMN, GH, JM, TTY, and EG contributed code to the vg toolkit (*annotate, pack, call* subcommands) with support from BP. AMN integrated new code into HTSlib. AMN and DC implemented new features in the Sequence Tube Map with support from GN. SD selected and analyzed the ATAC-seq datasets from ENCODE. JM conceived the study, analyzed data, and drafted the manuscript. All authors contributed to reviewing the text and content of the manuscript.

## Supplementary material

